# Integrative networks illuminate biological factors underlying gene-disease associations

**DOI:** 10.1101/062695

**Authors:** Arjun Krishnan, Jaclyn N. Taroni, Casey S. Greene

## Abstract

Integrative networks combine multiple layers of biological data into a model of how genes work together to carry out cellular processes. Such networks become more valuable as they become more context specific, for example, by capturing how genes work together in a certain tissue or cell type. We discuss the applications of these networks to the study of human disease. Once constructed, these networks provide the means to identify broad biological patterns underlying genes associated with complex traits and diseases. We cover the different types of integrative networks that currently exist and how such networks that encompass multiple biological layers are constructed. We highlight how specificity can be incorporated into the reconstruction of different types of biomolecular interactions between genes, using tissue-specificity as a motivating example. We discuss examples of cases where networks have been applied to study human diseases and opportunities for new applications. Integrative networks with specificity to tissue or other biological features provide new capabilities to researchers engaged in the study of human disease. We expect improved data and algorithms to continue to improve such networks, allowing them to provide more detailed and mechanistic predictions into the context-specific genetic etiology of common diseases

## B. INTRODUCTION

Many biological concepts and their interactions can be represented as networks. Existing types of networks used in computational analyses include networks of correlation based on gene expression [1, 2], protein-protein interactions [3, 4], and even phenotype-phenotype interactions [5, 6]. Some networks [7, 8] incorporate data from multiple layers of regulation including the interactome, transcriptome, and proteome.

In addition to data integration, a significant frontier for biological networks has been to include various forms of context specificity such as capturing interactions related to a certain process [9, 10] in a specific tissue [11, 12, 8]. This specificity has been achieved by either overlaying multiple resources of interactions [11] or employing a machine learning process that captures context specificity [8].

Biological networks, once constructed, provide a unique resource for the study of human disease. In many cases, phenotypes are expected to arise due to changes in how information flows through a biological system. For example, somatic mutation events in pancreatic tissue that constitutively activate KRas provide a constitutive growth signal to the system that can result in uncontrolled proliferation and pancreatic ductal adenocarcinoma [13, 14]. Large-scale biological networks serve as systems-level molecular scaffolds of cells on which researchers can locate known disease-associated genes, interpret their relationships in the context of other genes, and gain insights into how these genes might be involved in the disease. Given that the genetic bases of most complex diseases are poorly characterized, these networks also provide a genomic framework for identifying novel genes linked to diseases based on their patterns of network connectivity. These ‘interpretive’ and ‘predictive’ modes are often used in tandem with one cyclically feeding the other towards filling gaps in our knowledge of disease biology.

## C. INTEGRATED AND MULTI-OMIC NETWORKS

Biological networks can be constructed in many different ways and from many types of data. Correlation networks, for example, are one of the first large-scale models of gene interactions built solely based on patterns of shared gene expression and can be used to suggest opportunities for drug development or repurposing [2]. These can be global or tissue-specific [15]. Integrative, also called multi-omic, approaches, on the other hand, combine data-types across levels of biological regulation. One way to combine multiple data types is to overlay distinct information from separate data types. Okada et al. [16] combined data from text mining, risk-associated variants, protein-protein networks, molecular pathways, mouse phenotypes, and other sources to identify biological support for potential rheumatoid arthritis drugs. Such approaches look for broad support for the captured interactions based on evidence in all (intersection) or at least one (union) type of data without explicitly modeling distinct layers of data together.

Approaches that model multiple data types vary in how much they condition on known biological processes. Those that use documented regulatory patterns, e.g. one protein inhibits transcription of another gene, can reach a high level of detail in the constructed models. PARADIGM is an example of this type of approach. The method uses biological regulatory patterns to integrate multi-omic cancer measurements into a model that estimates the activity of individual proteins or pathways in a given tumor biopsy [17]. PARADIGM takes advantage of multiple data layers for the same samples. This makes it and methods like it well suited to the analysis of cancer genomics data where such opportunities are plentiful, but such methods are not as well suited to the analysis of large public data compendia. In these collections such matched data is often unavailable.

Methods that integrate public data into networks can first transform these datasets into scores for each gene pair. Some data types, such as protein-protein interactions, are already pairwise: edges exist for pairs of genes that encode proteins that physically interact. Other data types must be wrangled into pairwise relationships. For example, genome-wide expression information can be converted to correlations between each gene pair. Once converted to pairwise scores, machine learning methods allow researchers to combine different datasets, including gene expression and protein-protein interaction information, into a single network [18, 19, 10, 9]. These integrated networks capture broad biological processes and can be used for gene function prediction or other tasks [20, 18, 21]. In this review, we discuss their applications that aid in understanding the genetic and genomic basis of human phenotypes.

## D. INCORPORATING TISSUE-SPECIFICITY INTO NETWORKS

Constructing the biological gene networks specific to the hundreds of tissues and cell types in multicellular organisms is a major goal towards applying network biology to higher organisms. Pursuing this goal is to realize Waddington’s vision of “the complex system of interactions” – pegs representing genes and strings representing their “chemical tendencies” – underlying developmental landscape pulled by interactions anchored to genes [22]. This goal is being actively pursued using a variety of approaches addressing different facets of the challenge such as capturing specific types of gene interactions (e.g., functional, physical, or regulatory) and expanding the coverage of these networks to all genes in the genome and all tissues and cell-types in the body.

Overlaying gene (co-)expression, obtained from samples of a particular tissue, on protein-protein interaction (PPI) networks has been a straightforward and popular way to generate tissue-specific molecular networks [23]. Magger et al. [11] have shown that tissue-specific PPI constructed in this manner are valuable for disease-gene prediction using label-propagation methods. Comprehensive curation and comparison of multiple sources of tissue-gene expression has since been carried out [24, 25], results from which can be used to create tissues-specific PPI for an expanded set of genes and tissues. As an example, Cornish et al. [26] have incorporated more tissues and cell types to show that how tightly interconnected disease genes are in such tissue-specific networks can reveal interesting links between diseases and tissues. While comparing favorably with known associations from the literature, their results also point to novel tissue-disease links including one implicating mast cells in multiple sclerosis. These ‘data-overlay’ approaches succeed when many high quality documented annotations of gene expression and interaction are available.

Tissue-specific networks can also be generated using tissue-specific genome-wide data using approaches that do not require numerous documented annotations. Regulatory networks with tissue-specificity have been constructed by mapping transcription-factor binding sites to open chromatin regions in different tissues/cell-types identified based on DNase I sensitivity [27]. Alternative efforts have inferred networks by employing tissue-expressed promoter and enhancer elements [28]. Still others have developed inference methods to mine functional [29] or regulatory relationships [30] from large amounts of gene-expression data from a single tissue. Such approaches are particularly well suited when the cells being assayed (on a large-scale) most closely reflect the situation in complex human tissues.

However, large troves of existing genomic data available in public repositories, especially hundreds of thousands of gene-expression samples, are not resolved to tissues and cell-types. This represents a three-fold problem: 1) Many datasets are not annotated to the tissue/cell-type of origin due to unclear or entirely missing metadata; 2) Most samples are cell-type/tissue mixtures; and 3) Many cell-types are hard to isolate at all or enough to profile gene-expression. This challenge hence requires alternative approaches for integration and network inference.

A third type of approach aimed at addressing this challenge relies on applying machine-learning methods to simultaneously extract tissue and functional signal from data compendia representing heterogeneous tissue collections. Networks generated by these methods aim to be complete and predictive in general, however specific edges may be difficult to interpret. Work in *Caenorhabditis elegans*[31] and *Mus musculus*[32] demonstrated the potential viability of these approaches. Recently, tissue-specific networks constructed in this manner were generated for humans as well [8]. To perform this analysis, the authors developed gold standards of expression for 144 human tissues. They combined this with a gold standard of relationships within cellular pathways to generate 144 tissue-specific gold standards. The machine-learning process shown in Figure 1 constructs tissue-specific models from more than 32,000 experiments, most of which were not annotated or resolved to specific tissues. The models were then used to produce tissue-specific networks for each tissue. Comparison of tissue-specific networks constructed in this manner to those constructed only on tissue-specific data revealed consistent improvements with the machine-learning based integration of the complete data compendium. In addition to improved performance, the machine-learning approach could also be applied to generate networks for more tissues than the integration restricted to tissue-specific data.

**Figure 1:**
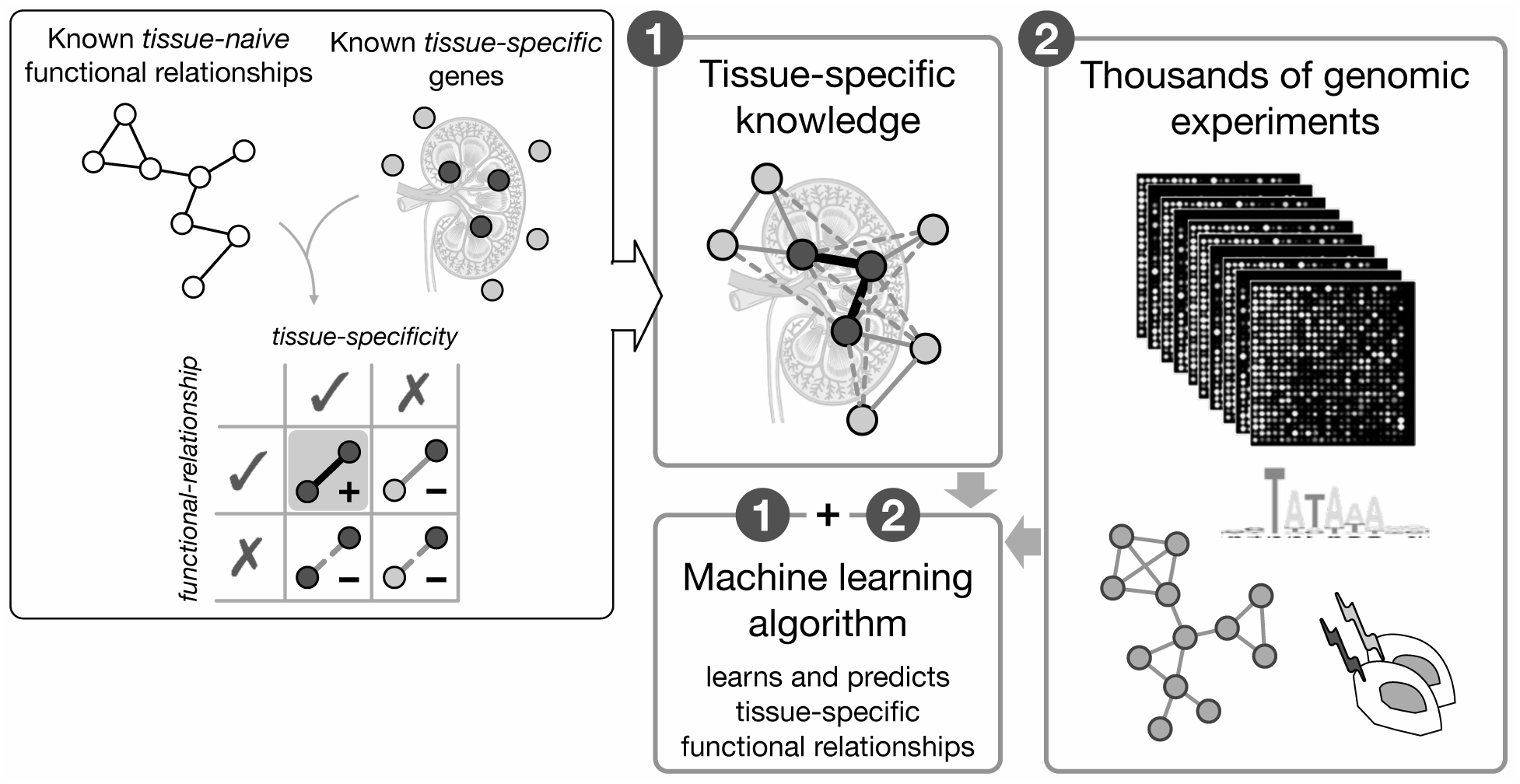
Integrating genomic data in the context of tissue- and functional knowledge to generate tissue-specific functional gene interaction networks. Nodes represent genes and edges represent specific relationships. Known tissue-naïve functional relationships (pathway/process co-membership links that are not specific to any tissue) are gathered from databases such as Gene Ontology. Known tissue-specific genes (black: genes expression in tissue of interest, say, kidney; grey: genes expressed in other unrelated tissue) are gathered from databases such as HPRD. Both tissue-naive relationships and tissue-specific genes are manually curated in these resources based on high-quality low-throughput experimental evidences. Combining the two produces a gold standard that represents our knowledge of ‘positive’ (both genes expressed and functionally related; thick black edges) and ‘negative’ (either gene in not expressed or genes not functionally related; full and dotted grey edges) gene interactions relevant to that tissue (labeled 1). Complementary to this low-throughput knowledge, thousands of genome-scale high-throughput experiments in the form of gene-expression profiles, protein physical interactions, genetic perturbations, and regulatory sequence patterns are available from public databases (labeled 2). A machine-learning algorithm can integrate all these of genomic datasets weighted by their relevance for and accuracy in capturing the tissue-specific knowledge of a given tissue. The algorithm learns distinctive patterns in genomic data that are characteristic of positive interactions and used these patterns to then predict the tissue-specific functional relationships between all pairs of genes in the human genome.

## E. APPLYING TISSUE-SPECIFIC NETWORKS TO STUDY HUMAN DISEASE

Human diseases are complex, and it is now clear that in many cases weak association signals are revealing broad networks of variants associated with such phenotypes [8, 33]. These may be numerous variants of weak effect acting additively, or the single-variant projections of complex epistatic disease models [34, 35]. In either case, the challenge now is to identify the biological signal hidden in a noisy set of statistical associations [36].

Networks provide a means to investigate these associations and potentially separate the disease-associated signal from statistical noise. Network-based approaches to identify and interpret disease associations are demonstrating success across diverse diseases including coronary artery disease [37], hypertension [8], cancer [38–40], multiple sclerosis [41], Alzheimer’s disease [42], and autism spectrum disorder [43]. Network-based approaches also present unique challenges – e.g. genes with more single nucleotide polymorphisms (SNPs) are also more likely to be highly connected within the network [44]. Approaches that employ network-based analysis should carefully evaluate and control for such biases.

In addition to considering networks, tissue-specificity of either causes or symptoms is a key feature of many diseases, and work by Lage et al. demonstrated the importance of considering tissues [45]. The importance of tissue-specificity has carried over to recent work that examines many genetic associations for a specific disease, For example, rheumatoid arthritis is an autoimmune disease. Walsh et al. [46] recently demonstrated that genetic associations with the disease reveal how those variants can impact cell-lineage-specific regulation to contribute to the etiology of the disease. There are a number of strategies for incorporating tissue-specificity into the network analysis of disease. We review four approaches that use network analysis of associations of SNPs, gene expression, or exome-sequencing to identify factors underlying complex diseases.

Tissue-specific networks can identify biologic commonalities underlying variants associated with human phenotypes and use those commonalities to identify disease-associated genes by their network connectivity patterns. In Greene et al. [8] the authors constructed 144 tissue-specific networks of predicted intra-pathway relationships (Figure 1). They then developed a procedure called the network-wide association study (NetWAS). The NetWAS trains a classifier to identify genes that should be associated with a disease based on their network connectivity patterns. Specifically, genes with a nominal association are used as positives for a machine learning algorithm and those without a nominal association are used as negatives. A classifier is then trained with these labels and all pairwise network edges as features. The classifier is then applied back to the network to identify genes with edges that indicate potential associations. This approach outperformed a hypertension genome-wide association study (GWAS) alone, and the network-prioritization process also ranked the gene targets of antihypertensive drugs more highly than GWAS. This suggests that association-guided network-based approaches may also aid drug development and repurposing.

Tissue- and cell type-specific networks that integrate a large amount of data can be particularly valuable for the study of rare diseases. For instance, systemic sclerosis (SSc) is a rare autoimmune disease characterized by fibrosis in skin and internal organs (e.g., pulmonary fibrosis). The challenges of studying of SSc at the molecular level are common to many rare diseases: sample sizes tend to be small and internal organ biopsies are often difficult to obtain. Thus, there is a critical need for leveraging additional biological data to make inferences about rare disease pathobiology. Taroni et al. [47] mined gene expression spanning multiple tissues and clinical manifestations in SSc to derive a disease associated gene set used to query the tissue-specific networks from Greene et al. [8]. The authors then performed differential network analysis on skin- and lung-specific networks to compare disease signal in the major organ systems affected by fibrosis. The authors first identified gene-gene interactions that were highly specific to each tissue by subtracting the global edge weights from the skin and lung networks. Then, these highly tissue-specific edges were compared to identify functional differences. They found a set of edges highly specific to lung and suggestive of a distinct macrophage phenotype in lung as compared to what could be inferred from SSc skin gene expression data. The authors also developed a cell type-aware multi-network approach that detected genes preferentially downregulated in skin during improvement of immunomodulatory treatment. This work suggests tissue- and cell type-specific functional genomic networks can provide insight into rare disease processes that are difficult to capture experimentally.

The advent of high-throughput sequencing has enabled the discovery of regulatory variants relevant to disease especially through studying the genetics of gene/protein expression [48]. Projects such as the GTEx [49, 50] and FANTOM5 [51] have used this approach to highlight regulatory variants that are specific to or shared across tissues. Analogous to tissue-specific functional networks, tissue-specific regulatory networks that can be inferred from such data have the potential to delineate the role of regulatory variants in specific tissues. Marbach and colleagues [28] inferred tissue-specific regulatory networks by overlaying transcription factor binding-site occurrences over promoter and enhancer elements detected in hundreds of tissues/cell-types in the FANTOM5 project. They then use this network to assess links between diseases and tissues by calculating how surprisingly tightly connected are disease-genes implicated by GWAS in a particular tissue regulatory network (termed connectivity enrichment analysis). This analysis entails ranking genes by their summarized GWAS p-values, calculating the average connectivity of the genes above each rank along the list, and estimating the area under the curve (AUC) of connectivity as a function of rank. A connectivity score is finally reported based on the comparison of the observed AUC with an empirical distribution of AUCs generated by performing the analysis on thousands of permuted rankings. By performing this analysis across multiple diseases and networks, the authors find that disease genes are indeed tightly clustered in tissues relevant to the disease.

Traditional network-based disease-gene prediction algorithms take as input either only known high-confidence disease-genes or all genes irrespective of evidence. Krishnan and colleagues [43] have recently developed an evidence-weighted approach that incorporates the trust with which disease genes are known from the sources of evidence be it high-confidence candidates identified in sequencing studies or weak associations mined from literature. They used this approach in conjunction with the brain-specific network, developed previously [8], to discover novel candidate genes associated with autism spectrum disorder (ASD). Intuitively, the underlying machine learning method learns the pattern of network connectivity characteristic of known ASD genes (weighted proportional to their evidence) and then identifies other genes in the network that highly resemble known genes in their network pattern. The method is highly general and can be applied to any complex disease to predict new genes based all the genes previously identified in single exome/whole-genome sequencing studies or collated from multiple datasets.

Taken together, integrative networks serve as powerful means to both interpret existing knowledge about complex physiology and disease, while also offering data-driven predictions that point to uncharted territory, be it novel genes associated with diseases, new pathway memberships, key regulators of disease genes/pathways, surprising associations between tissues and diseases, or differential effects of disease on different tissues. Based on their interest and scope, experimental/biomedical researchers use these network-based tools to identify a small number of targets for careful investigation or to gather a prioritized list of candidates to guide further large-scale genetic screens.

## F. THE ROAD AHEAD

A major bottleneck in building accurate tissue- and cell-type-specific networks is the extreme scarcity of prior knowledge about tissue-specific genes and interactions based on high-quality experimental evidence. As we amass even a few scores of such specific genes and interactions across under-studied tissues/cell-types, they can be used to train better machine-learning classifiers and serve as reliable evaluation standards. A lesser but still significant challenge is the unavailability of large-scale genomic data uniformly from normal and disease states across a wide range of tissues/cell-types. Such biases in input datasets, along with scarce prior knowledge, creep into the final inferred networks for different tissues/cell-types, affecting their robustness and genomic coverage. In contrast, rare examples of massive efforts focused on a single tissue/cell-type help us describe the scope of data needed to construct better networks. Such efforts are fueled by a large collection of data from a single tissue/cell-type including transcriptional profiles across a range of different genetic perturbations and time points, sometimes further profiling protein-protein or protein-DNA binding on a large-scale, which together facilitate integrative methods to reconstruct accurate networks [52, 53]. Overall, both approaches broadly covering many tissues and deeply covering single cell-types have enormous merits to be pursued simultaneously and, ideally, feed into each other.

If sufficient gene expression data is available at least for multiple tissues in healthy and a particular disease condition, it may be possible to utilize differential network analysis to study disease-specific processes across tissues akin to [47]. We illustrate with a hypothetical example centered on blood and a disease end-target tissue, kidney, as an example for how this strategy may be used to gain insight into disease mechanisms. Tissue- and condition-specific networks could be learned from four compendia of data (disease blood, healthy blood, disease kidney, and healthy kidney) using the same functional standards and then compared. If an intra-pathway relationship exists in both disease and healthy blood, but that intra-pathway relationship is absent in healthy kidney when compared to disease, we might infer that a leukocyte population is present in the diseased tissues. Furthermore, if the intra-pathway relationships identified in blood were connected to other pathways in diseased tissues (blood and kidney) alone that would suggest immune infiltrate with a particular phenotype in diseased kidney. With data available on genetic variants, it could also be possible to link cell-type specific changes in networks to underlying causal variants [54]. These multi-tissue differential network analyses could then guide further analyses and experiments centered on this complex multi-cell lineage disease process.

In addition to the gene networks specific to individual tissues, another milestone for the biological networks is the ability to capture cross-tissue interactions. Much of human physiology relies on biochemical interactions such as hormonal signaling and immune response that span across tissues. Capturing these interactions requires particular data from multiple tissues of matched individuals. Dobrin et al. [55], proposed one of the earliest approaches towards this goal by calculating gene-gene correlation across hypothalamus, liver and adipose tissues in mouse and using the resulting networks to study molecular crosstalk between these tissues related to obesity. However, coordinated changes in gene expression across tissues can arise due to a common underlying genetic or regulatory mechanism. Such mechanisms could induce changes in both tissues, without reflecting bona-fide cross-tissue interactions. The recently produced large multi-tissue, multi-individual gene expression data from the GTEx consortium helps to address this challenge. Long and colleagues [56] calculated inter-tissue interactions while carefully controlling for the identical genetic regulation. Using these inter-tissue networks, they highlight specific signaling links between heart and whole-blood or lung. Similar to the methods described above for constructing tissue-specific networks, another recent study [57] has taken a network-overlay approach to infer cross-tissue interactions. In this study, Ramilowski and colleagues overlay tissue/cell-type gene-expression data from the FANTOM5 database onto known physical interactions between ligands and receptors, identifying cases where the ligand is expressed in one tissue and the receptor in another. While these methods are beginning to shine light on important aspects of human biology, the ground is fertile for novel integrative methods that can construct such cross-tissue maps on a large scale.

As network-based approaches are gaining use in genetic association analysis of individual complex diseases, another natural leap is to the domain of multiple-phenotype analyses. Because many genes and pathways have pleiotropic effects, i.e. their activity can alter multiple phenotypes, phenome-wide association studies (PheWAS) allow for this additional information to be incorporated. Complementary to identifying multiple variants associated with a trait of interest using GWAS, PheWAS interrogates the association of a variant of interest with a range of traits/phenotypes. Phenotype data for such an analysis are abundant, albeit in a noisy manner, in electronic health records (EHRs) of thousands of patients. If these patients have also been genotyped, PheWAS can be used to link variation at those locations to EHR-derived phenotypes or clinical outcomes that vary in the patient population [58]. Denny and colleagues [59] have applied this approach to specifically discover several novel pleiotropic variants associated with multiple phenotypes, a feat that is challenging in a disease-centric setting as in GWAS. Although a few previous studies have used a disease network (inferred based on phenotypic similarity) in combination with a gene network to model many-to-many disease-gene associations [60], much work is needed to develop network-based approaches that can complement PheWAS in identifying the tissue-specific effects of pleiotropic genes.

Most networks approaches developed to date have dealt with a single data type or a single integrated portrait from multiple datatypes over a common biological entity, the gene. New methods are being developed that incorporate multiple network types into a single heterogeneous network (‘hetnet’) [41]. These methods have helped to prioritize disease-associated genes [41], and the ongoing Project Rephetio [61] provides a promising method for drug repurposing using these data. Though hetnets are in their early days, they may provide a powerful means to develop algorithms that can consider multiple biological entities simultaneously to identify the basis of human phenotypes.

## G. CONCLUSIONS

Progress to date on network-based methods to identify the basis of human phenotypes has been promising. The initial step: connecting genetic variants to the gene that they affect remains challenging [62]. Advances in this domain will naturally translate to improvements for network-based methods generally focus on genes. In addition to the supervised methods to construct networks highlighted in this review, new and powerful unsupervised learning approaches may allow us to construct networks in cases where biological knowledge is limited or unavailable [63]. While we anticipate that large-scale integrated networks will make their first contributions at the level of identifying a shared genetic or pathway basis behind observed associations, we look forward to detailed networks that can suggest a mechanistic hypothesis. These networks will require the combination of new algorithms and analytical methods as well as detailed data in targeted domains. Though more remains to be done, progress in this area is encouraging and we look forward to advances in the years to come.

Papers of particular interest, published recently, have been highlighted as:

* Of importance.

